# Functional diversity reduces the risk of hydraulic failure in tree mixtures through hydraulic disconnection

**DOI:** 10.1101/2023.06.09.544345

**Authors:** Myriam Moreno, Guillaume Simioni, Hervé Cochard, Claude Doussan, Joannès Guillemot, Renaud Decarsin, Pilar Fernandez, Jean-Luc Dupuy, Santiago Trueba, François Pimont, Julien Ruffault, Nicolas K. Martin-StPaul

**Author notes:** (corresponding author) Myriam Moreno. **Email:**.

## Abstract

Forest ecosystems are increasingly threatened by anthropogenic pressures, especially by the increase in drought frequency and intensity. Tree species mixtures could improve resilience to diverse global anthropogenic pressures. However, there is still little consensus on how tree diversity affects water stress. Although some studies suggest that mixing species with different drought response strategies could be beneficial, the underlying mechanisms have seldom been identified. By combining a greenhouse experiment and a soil-plant-atmosphere hydraulic model, we explored whether mixing a drought avoidant (*Pinus halepensis*) and a drought tolerant (*Quercus ilex*) tree species could reduce plant water stress (defined as the risk of hydraulic failure) during extreme drought, compared to their respective monocultures. Our experiment showed that mixing species with divergent drought response strategies had a neutral effect on the drought-avoidant species and a positive effect on the drought-tolerant species. The model simulations further suggested that the beneficial effect of mixture on plant water stress during extreme drought was related to changes in the hydraulic connection of the plant from both the soil and the atmosphere. The ability of the drought-avoidant species to disconnect from the soil and the atmosphere limits its exposure to water stress, whereas the ability of the drought-tolerant species to increase its hydraulic connection to the soil lowers its hydraulic risk. This study brings a new insight on the mechanisms and traits combinations improving drought resistance in diversified forests and plantations, with important implications for forest management under climate change.

## Introduction

The rising frequency and intensity of extreme drought is impacting tree survival and forest functions worldwide (Allen et al., 2010; Breshears et al., 2013; Senf et al., 2020), jeopardizing crucial forest ecosystem services. Tree species diversity has been promoted as an important nature-based solution to improve the resilience of forests and tree plantations (Messier et al., 2022). The effects of species mixing on drought resistance could result from different mechanisms, such as competitive reduction for water through resource partitioning or facilitation – for instance hydraulic redistribution (Grossiord, 2020). Yet, there is no consensus regarding the effects of tree diversity on forest resistance to drought (Grossiord et al., 2014; Grossiord, 2020). Indeed, recent review showed that diversity can have positive (de-Dios-García et al., 2015; Lebourgeois et al., 2013; Ruiz-Benito et al., 2017), neutral (Grossiord et al., 2014; Merlin et al., 2015) or even negative impacts (C. Grossiord et al., 2014; Vitali et al., 2018). These conflicting results suggest that it is not the species richness that matters, but rather the functional composition (i.e., species with different drought response strategies) of the mixtures (Forrester and Bauhus, 2016; Grossiord, 2020). Such hypothesis was supported by recent research that found that the diversity of hydraulic traits determines the resilience to drought of forest water fluxes globally (Anderegg et al., 2018; Haberstroh and Werner, 2022). Similarly, results from a large-scale tree diversity experiment showed that the diversity of drought resistance strategies is a good predictor of the stability of tree growth and forest productivity (Schnabel et al., 2021). However, we crucially miss a mechanistic understanding of the way the diversity of drought resistance strategies mediates tree mortality under extreme drought.

Tree drought resistance strategies result from a set of functional traits that determine how rapidly the different tree functions will be impaired by drought stress (often quantified as water potential thresholds inducing dysfunction). In particular, it determines the risk of xylem hydraulic failure, caused by a high rate of embolism in xylem conduits (Tyree and Sperry, 1989), which is a leading mechanism in drought-induced tree mortality (Adams et al., 2017). It is common in the literature to distinguish species strategies based on stomatal regulation - and associated water potential dynamics - and the xylem vulnerability to embolism (Chen et al., 2021). Drought-tolerant species tend to maintain gas exchanges during drought by delaying stomatal regulation, which implies important soil water depletion and large decrease in the soil and tree water potential during drought (Delzon, 2015; López et al., 2021). Their high resistance to xylem embolism limits the risk of hydraulic failure. By contrast, drought-avoidants are generally more vulnerable to xylem embolism, but they close their stomata earlier during drought, thereby reducing soil water depletion, which in turn limits the soil and tree water potential decrease and the risk of hydraulic failure (Delzon, 2015; López et al., 2021).

By assuming that trees are hydraulically connected to the soil (*i.e.,* soil and tree water potential are at equilibrium if transpiration is null such asunder predawn conditions) and that the root system fully occupies a given soil volume, one can hypothesize how mixing species with distinct drought response strategies impacts soil and tree water potentials, and the risk of hydraulic failure under extreme drought:

**H1.** For a drought-tolerant species, it should be beneficial to compete for water with a drought-avoidant neighbour, because the soil water saved by earlier stomatal regulation of the avoidant is available to delay the decrease in water potential and the overall hydraulic failure risk (Fig. 1).

**Figure 1.**
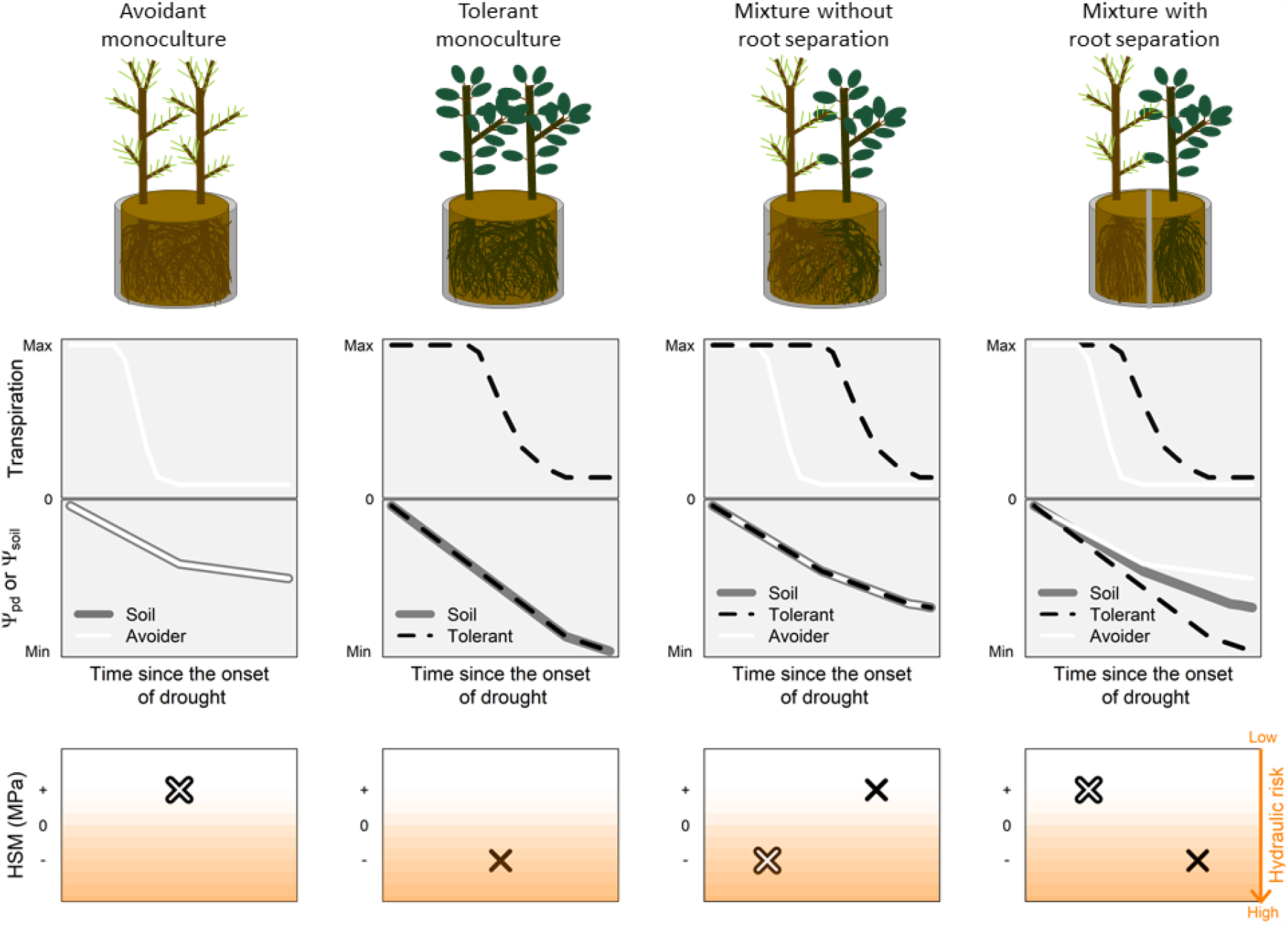
Experimental design and hypothesized drought responses for monoculture and mixture of a drought avoidant and a drought tolerant species. The transpiration, water potentials (Ψ_soil_: overall pot soil water potential; Ψ_pd_: tree water potential) and hydraulic safety margins (ie. HSM: the difference between Ψ_pd_ and P50; the water potential causing 50 % of embolism) expected according to pot modalities are presented. In the drought avoidant monoculture, trees transpiration are expected to reduce rapidly after the onset of drought, limiting Ψ_soil_ and Ψ_pd_ drop and hence hydraulic failure risk (positive HSM). In the drought tolerant monoculture, transpiration of the two trees is expected to reduce later than the one of the drought avoidant species, inducing a sharp decrease of Ψ_soil_ and Ψ_pd_, increasing the risk of hydraulic failure (negative HSM). In the mixture without root separation, transpiration of the drought avoidant species decreases earlier than for the drought tolerant, which improve the water potential and HSM of the drought tolerant compared to monoculture. However, because the two trees share the same volume of soil, the water consumption of the drought tolerant should decrease water potential and HSM of the drought avoidant thereby increasing its hydraulic failure risk compared to the monoculture. A mixture with root separation illustrates that when each species root system occupies its proper soil volume, the regulation of the transpiration, the water potentials dynamics and the HSM are expected to be the same as in monoculture. As Ψ_soil_ represents the global pot soil water potential, it is here equal to the mean of both compartment soil water potential.

**H2.** When grown in mixture with a drought tolerant neighbour, a drought-avoidant species should be disadvantaged, as it would experience lower soil water potential due to sustained water-use by the companion tolerant species. This would lead to a decrease in its water potential, thereby increasing the risk of hydraulic failure (Fig. 1). The scenario presented in Fig. 1 - which suggests that a drought tolerant always “win the fight” during drought under mixture - holds only if the water potential of the mixed species is at equilibrium with the soil water potential.

**H3.** If the root systems of the two neighbour species are segregated in space, water consumption by the tolerant species does not affect the avoidant species and difference of water potentials between tree species in the mixture could occur given their isolation (Fig. 1). In support to this hypothesis, root niche separation is often assumed in the literature to explain coexistence between co-occurring species (Grossiord, 2020; Jose et al., 2006).

To test these hypotheses, we conducted a greenhouse experiment where seedlings of *Pinus halepensis*, a drought avoidant (Baquedano and Castillo, 2007) and *Quercus ilex*, a drought tolerant (Baquedano and Castillo, 2007) were grown in pairs in small pots (12 L). Seedlings were planted in either monoculture or mixture with or without root separation. We applied an extreme drought by stopping the watering and we regularly monitored the overall pot soil (Ψ_soil_) and tree predawn (Ψ_pd_) water potentials, along with soil resistivity and tree gas exchanges. These data were combined with a state-of-the-art soil-tree-atmosphere hydraulic model (Cochard et al., 2021) to further identify the mechanisms and traits involved in the co-existence of drought avoidant/tolerant species during extreme drought.

## Results and Discussion

### a) Empirical evidence for hydraulic commensality between *Quercus ilex* and *Pinus halepensis* during extreme drought

Tree predawn water potential (Ψ_pd_) decreased markedly during drought (SI appendix, Table S1, P-value < 0.001), but the dynamics differed between species across treatments (SI appendix, Table S1, P-value < 0.001). Due to their differential drought resistance strategies, Ψ_pd_ reached more negative values in the drought tolerant *Q. ilex* than in the drought avoidant *P. halepensis*, regardless of the pot composition. In agreement with hypothesis H1, *Q. ilex* had significantly more negative Ψ_pd_ in monoculture than in mixture at the drought peak (Fig. 2A; SI appendix, Table S2, P-value < 0.01), except for the mixture with root separation. By contrast, in *P. halepensis*, Ψ_pd_ at the drought peak remained above both soil water potential (Ψ_soil_) and *Q. ilex*’s Ψ_pd_ in mixture (Fig. 2A), contradicting hypothesis H2.

**Figure 2.**
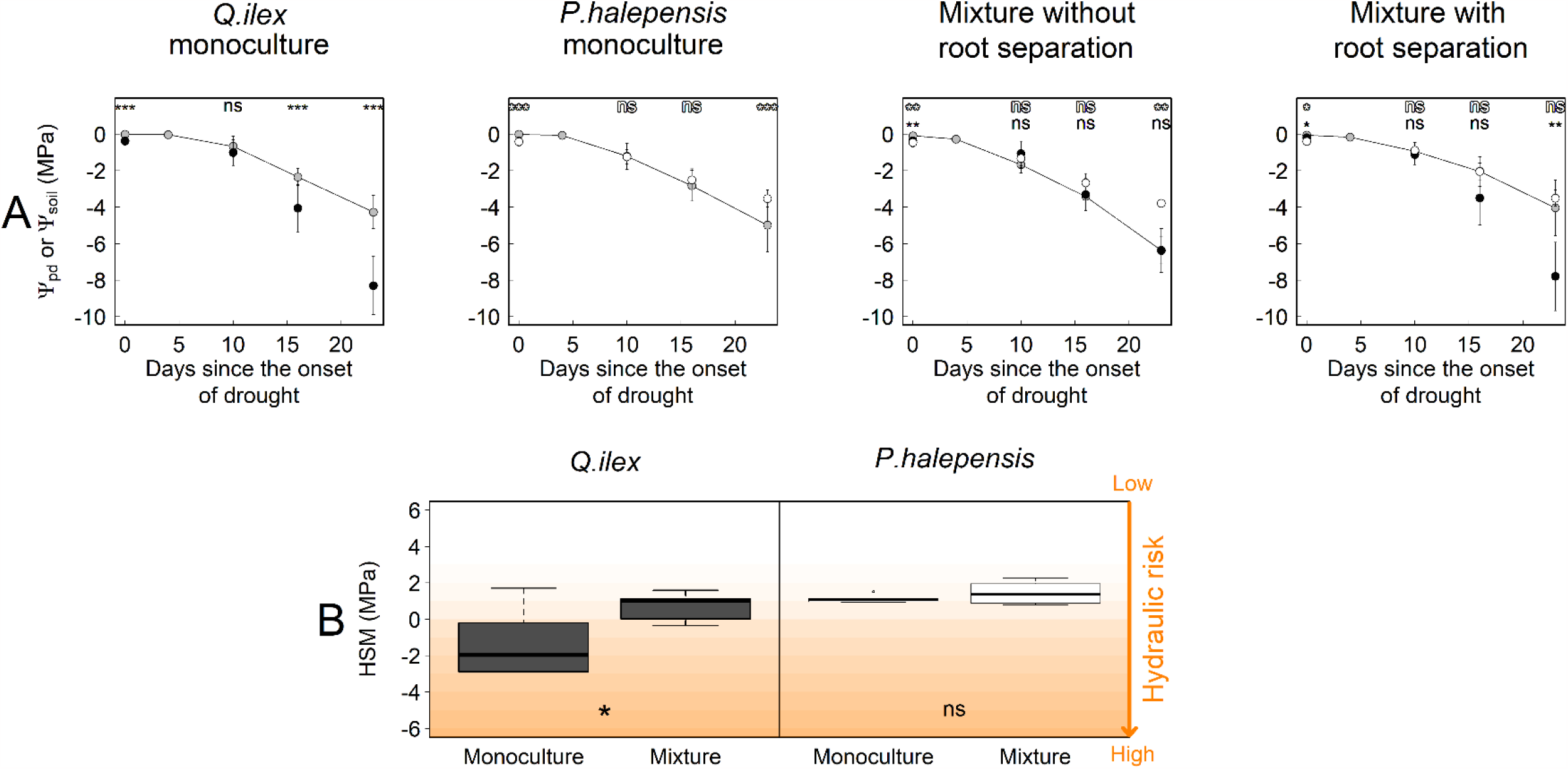
Positive and neutral effect of mixture on hydraulic failure risk of drought tolerant *Q. ilex* and drought avoidant *P. halepensis*. (A) Soil and tree water potential for the different pot composition at each measurement dates. Soil water potentials represent average values computed at the pot level from manual weightings (grey points). The average tree water potentials of *Q. ilex* and *P. halepensis* correspond respectively to black and white dots. Standard deviations are represented and significant differences between soil and water potentials are indicated (ns, non-significant differences; *, 0.01 ≤ p < 0.05; **, 0.001 ≤ p < 0.01; ***, p < 0.001). Per measurement date, for Ψ_pd_, N = 24 for monoculture (pooling monoculture with and without root separation/ two trees per pots) and 6 for mixture. For Ψ_soil_, N = 12 for monoculture (pooling monoculture with and without root separation) and 6 for mixtures concerning soil water potentials. (B) Hydraulic safety margins (HSM) measured at the driest date of the experiment in monoculture (with and without root separation) and mixture (only for pots designed without root separation). HSM were computed as the difference between water potential at the driest date and the P50 (i.e., the water potential causing 50% embolism).

Therefore, the hydraulic risk, estimated as the hydraulic safety margins at the drought peak (HSM, computed as the difference between P50, the water potential causing 50% xylem embolism and the average Ψ_pd_ at the driest date) was significantly improved in mixture compared to monoculture only for *Q. ilex* (Fig. 2B). Hence, as expected from hypothesis H1, the mixture had a positive effect on the reduction of the risk of hydraulic failure for the drought tolerant species *Q. ilex* but contrary to hypothesis H2, the mixture had a neutral effect on the drought avoidant *P. halepensis.* This result reflects a commensalism relationship in terms of hydraulic risk between drought avoidant and tolerant species under drought conditions, that to our knowledge, has never been demonstrated until now. Hence, coexistence between a drought avoidant and a drought tolerant species is not the exclusive result of spatial segregation of their root niche, even if such phenomenon can occur (Bello et al., 2019), but depend rather on other mechanisms that we further discuss below.

### b) Species coexistence relies on species-specific modifications of the soil-tree hydraulic conductance

The relationship between Ψ_pd_ and Ψ_soil_ (Fig. 3) in *P. halepensis* was unaffected by mixture, with Ψ_pd_ equal to Ψ_soil_ until Ψ_soil_ decreased below -4 MPa. For Ψ_soil_ lower than - 4MPa, Ψ_pd_ remained constant at ca. -4 MPa. For *Q. ilex* the slope between Ψ_pd_ and Ψ_soil_ was greater than one (> 1.7, Fig. 3) for monocultures and for the mixture with root separation. By contrast, the slope of the Ψ_pd_-Ψ_soil_ relationship was equal to one for *Q. ilex* in mixture without root separation (i.e., Ψ_pd_ ∼Ψ_soil_ throughout the experiment, Fig. 3). Overall, these empirical results indicate that (i) the Ψ_pd_ *vs*. Ψ_soil_ relationships varied between the two studied species with different drought resistance strategies and (ii) plant-soil water potentials were modified by mixture only in *Q. ilex*, the drought tolerant species.

**Figure 3.**
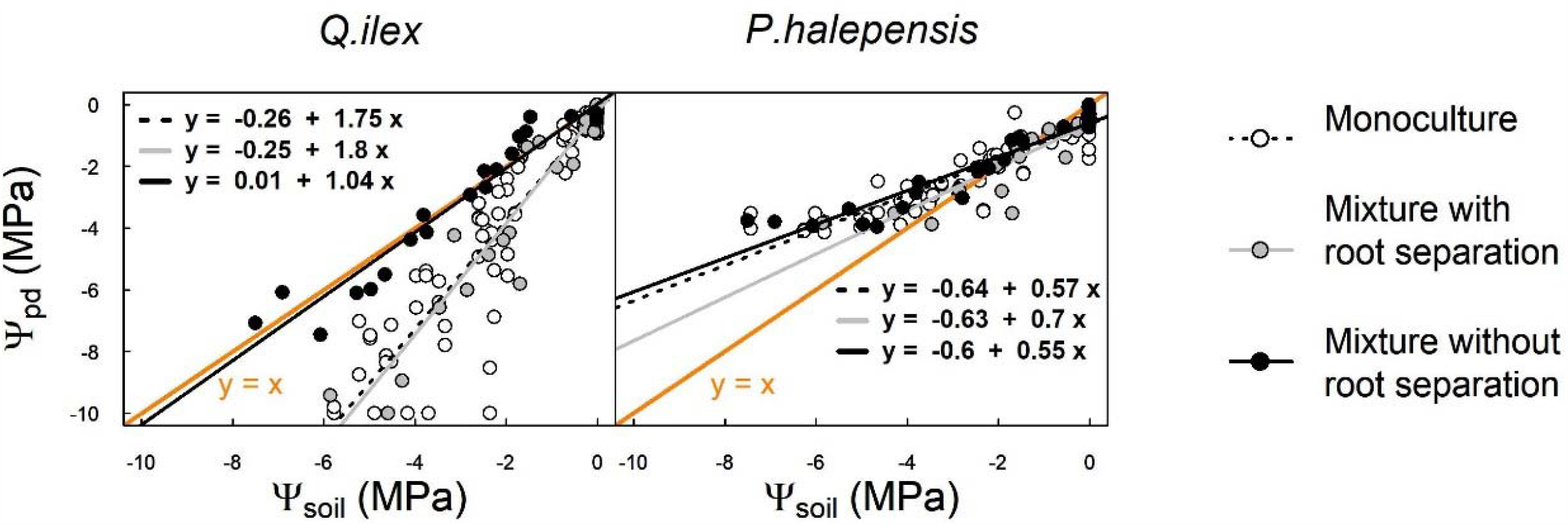
Uncoupling between soil and tree water potentials suggest an improvement of the soil-tree hydraulic conductance for *Q. ilex* in mixture and soil isolation for *P. halepensis*. Different colours were used for monoculture (black dots), mixture with root separation (light grey dots) and mixture without root separation (dark grey dots). The isoline (y=x) is reported in orange line. For each modality, linear fit between soil and water potentials is depicted and the equation is indicated on the plot. N = 96 for monoculture (with and without root separation) and 24 for mixture for each root separation categories.

It could be assumed that differences between Ψ_pd_ and Ψ_soil_ reflect shifts in the root profile in mixtures compared to monocultures. Indeed, if roots explore only a part of the available soil, Ψ_pd_ would equilibrate with this soil subspace, possibly differing from the overall Ψ_soil_ measured at plot level. However, we used small pots (12 L) to impose a complete occupation of the whole soil volume by the trees’ root system, making this assumption unlikely. Furthermore, the fact that we found no significant differences between the average soil resistivity at the top and bottom profiles of the pots for each modality suggests that water is absorbed uniformly throughout the soil and definitely rules out this hypothesis (SI appendix, Fig. S1). Alternatively, we can postulate that differences between Ψ_pd_ and Ψ_soil_ result from changes in the hydraulic conductance between the soil and the trees.

Following the experimental component of our study, we carried out simulations with the hydraulic process-based model SurEau (Cochard et al., 2021) to test the possible mechanisms that could explain such empirical patterns. The model computes the water fluxes along the soil-tree-atmosphere continuum by accounting for the different resistances of the soil, the symplasm and apoplasm of the root, trunk, branch and leaf, and calculate the water potential and the water content of the corresponding compartments. By considering xylem vulnerability to cavitation, the model can estimate the loss of hydraulic conductance of the tree xylem in relation to water potentials and predicts the death of the tree by hydraulic failure when 100 % loss of hydraulic conductivity is reached. The model was improved to represent two different individuals competing for a same amount of soil water (see Materials and Methods).

We first conducted three benchmark simulations corresponding to Figure 1 (monocultures of *Q. ilex* and *P. halepensis* and the mixture without root separation). In such simulations, we assumed that the hydraulic conductance of the rhizosphere (*K*_rhyzo_) and of the fine roots (*K*_*root*_) were the same for all species and pot compositions. More specifically, we applied the widespread “single root” approach that assumes that soil conductivity relates to the soil water content (van Genuchten, 1980) and is scaled up to the rhizosphere according to the length of fine roots per unit soil volume (Gardner, 1964; I. R. Cowan, 1965)

By doing so, the model predicted behaviours consistent with our initial hypotheses H1 and H2 (Fig. 1): the drop of Ψ_pd_ in *P. halepensis* and *Q. ilex* growing in mixture are respectively faster and slower than for the corresponding monocultures (Fig. 4A), indicating that mixture should have a negative effect on *P. halepensis* and a positive effect on *Q. ilex*, which contradict our results. We thus conducted sensitivity analyses on different traits to explore mechanisms explaining our empirical observations.

**Figure 4.**
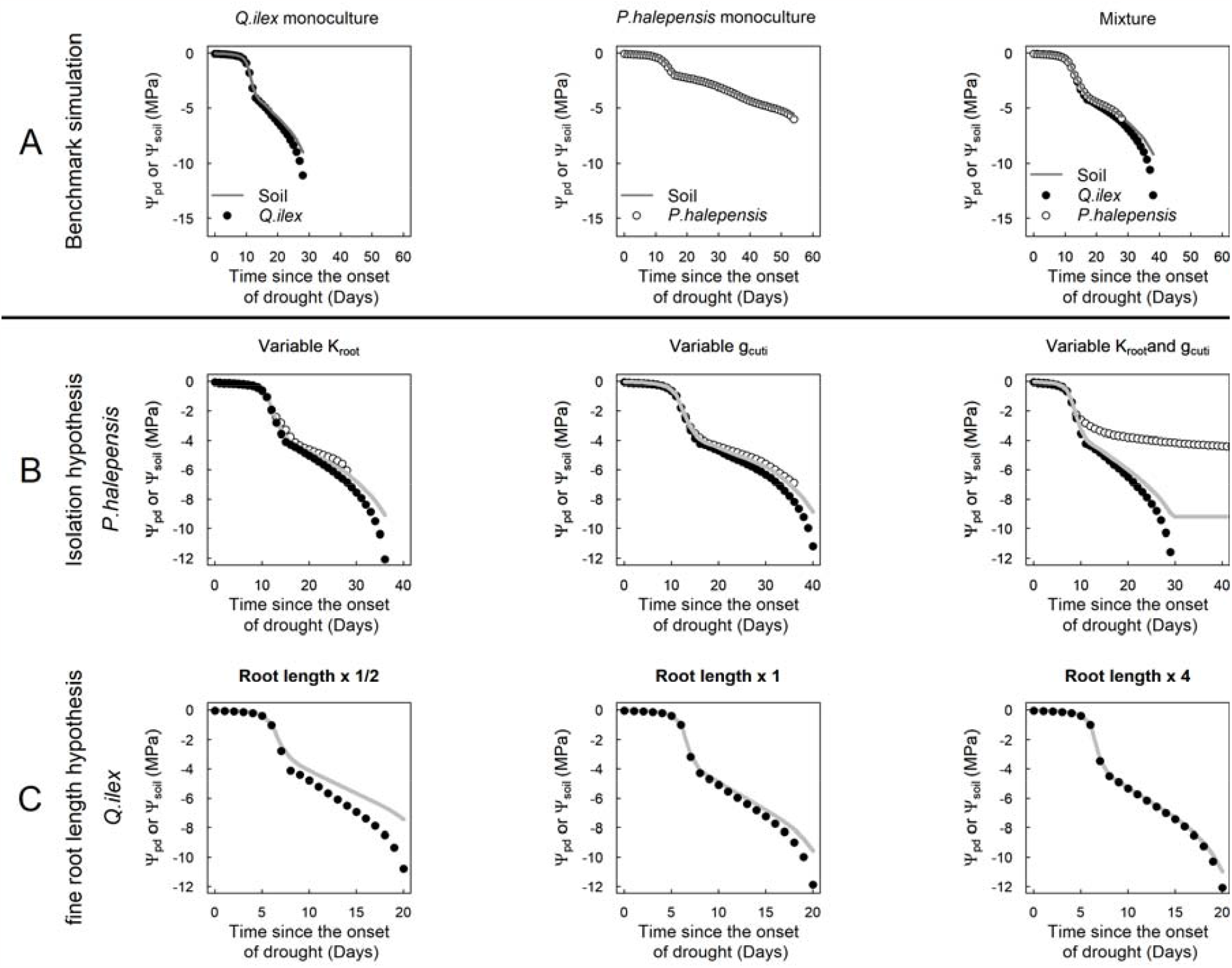
Implication of the soil-tree hydraulic conductance in the coexistence of drought avoidant and drought tolerant species during extreme drought. (A) Benchmark SurEau simulations of the dynamics of leaf and soil water potentials for *P. halepensis* and *Q. ilex* grown in monoculture and mixture until tree death. In these simulations root hydraulic conductance and cuticular conductance were kept constant. The results are in adequation with H1 and H2 hypotheses postulated in the introduction (illustrated in Fig.1). (B) Test of sensibility of root conductance (K_root_) and leaf cuticular conductance (g_cuti_) parameters for *P. halepensis*. By reducing both parameters, trees can keep higher water potentials than the soil. (C) Test of sensibility of fine roots length parameter for *Q. ilex* (multiplying fine roots length by ½, 1 and 4). The more the fine roots length, the closer are tree predawn and soil water potential. Note that the graduation of the x and y axis change according to plot. Model parameters are provided in the SI appendix, Table S4.

### c) The drought-avoider *P. halepensis* isolates from the soil through a decrease in both root hydraulic and cuticular conductance

The fact that *P. halepensis* exhibits higher Ψ_pd_ than Ψ_soil_ during drought suggests that (i) this species can isolate from the soil (i.e., reducing the soil to tree hydraulic conductance) and (ii) is able to limit its dehydration. Several studies have suggested that plant isolation from the soil allows limiting the exposure to water stress (Aguadé et al., 2015; Brito et al., 2019; Cuneo et al., 2016). Different non-exclusive belowground mechanisms were proposed to explain tree isolation from the soil, such as the formation of cortical lacunae under fine roots (Cuneo et al., 2021; Duddek et al., 2022), which reduces water transfer to the root stele and hence affect roots hydraulic conductance. Roots shrinkage might also explain the plant-soil hydraulic disconnection by creating gaps between soil and fine roots interrupting the hydraulic conductance between both interfaces. Furthermore, the inhibition of the synthesis of proteins such as aquaporins facilitating water transport in the transcellular pathway (Domec et al., 2021), or even fine roots mortality (Leonova et al., 2022). could also explain hydraulic isolation. To evaluate whether tree isolation from the soil could explain the observed water potential patterns in *P. halepensis*, we hypothesized in the model a decrease in root hydraulic conductance (*K*_*root*_) as the tree water potential decreases (SI appendix, Fig. S2). Simulations were performed under a mixture condition with *Q. ilex* as a companion species (parametrized as in benchmark simulations) (Fig. 4B). This allows to force soil water potential to drop even after *P. halepensis* has closed its stomata and has isolated from the soil. Model simulations indicate that reducing only *K*_*root*_ does not allow to simulate higher Ψ_pd_ than Ψ_soil_ for *P. halepensis* (Fig. 4B). This means that the water losses that occur after stomatal closure – which result from the leaf cuticular conductance (*g*_cuti_), set in the model using the average value measured for *P. halepensis*, was high enough to cause tree water potential drops after a strong decrease in *K*_*root*_ (Fig. 4B). We thus implemented in the model a down-regulation of *g*_cuti_ with decreasing tree relative water content, which is in accordance with empirical data obtained in *P. halepensis* using the drought-box methods (Billon et al., 2020) (SI appendix, Fig. S3). Simulations showed that, although the reduction of *g*_cuti_ alone attenuated the decrease in tree water potentials, the tree keeps dehydrating. Finally, when implementing a decrease of both *K*_*root*_ and *g*_cuti_ under drought, *P. halepensis* water potential departs from soil water potentials (Fig. 4B), in line with our observations. This suggests that these two mechanisms jointly could allow *P. halepensis* to prevent dehydration under drought. In the natural forest context, tree isolation from the soil during drought has already been proposed to explain the coexistence of drought-avoidant and drought-tolerant trees (Aguadé et al., 2015; Moreno et al., 2021; Pangle et al., 2012; Plaut et al., 2012). Yet, to our knowledge, the mechanisms leading to complete plant disconnection from the soil and the atmosphere had never been proposed until now.

### d) The drought-tolerant *Quercus ilex* increases root hydraulic conductance to the soil in mixture through increased root length

For *Q. ilex*, Ψ_pd_ was respectively lower or comparable to Ψ_soil_ under monoculture and mixture conditions (Fig. 2A) which suggests that (i) contrary to *P. halepensis, Q. ilex* is not able to limit its dehydration and (ii) the mixture likely impact the hydraulic conductance between the soil and the trees under drought.

According to the diffusion law, the lower departure between Ψ_pd_ and Ψ_soil_ that we observed for *Q. ilex* in mixture compared to monoculture, could result from an increase in the conductance of the rhizosphere, which could lower the water potential drops required for a given flux between the soil and the tree (39).

As we found a greater root system length in mixture than in monoculture (SI appendix, Fig. S4), we assumed that the increase in rhizosphere conductance might be achieved through an increase in exchange surface between soil and root (“single root” approach). We tested this hypothesis by varying the modelling parameters of fine roots length per unit soil volume. This sensitivity test shows that changing *K*_rhyzo_ can change the Ψ_pd_ *vs* Ψ_soil_ relationships between monoculture and mixture (Fig. 4C). Indeed, reducing the value of this parameter (graph “root length x ½”, Fig. 4C), results in a departure between Ψ_pd_ and Ψ_soil_ as observed in monoculture, whereas increasing it results in Ψ_pd_ and Ψ_soil_ being comparable, as observed in mixture. Interestingly, some studies have already reported modifications of the root system under mixture toward higher fine roots density (Sun et al., 2017; Wambsganss et al., 2021), identifying this phenomenon to a complementarity effect between species associated.

### e) Ecological implications

Our results provide evidence that mixing drought-avoidant and drought-tolerant species reduces the risk of hydraulic failure under extreme drought conditions at the community level. According to model simulations, such mixing effect can be explained by changes in hydraulic connection between the plant, the soil and the atmosphere during drought. The avoidant species can sustain extreme drought through an isolation from the soil (decrease of K_root_) and the atmosphere (decrease of *g*_cuti_) whereas the tolerant species can increase hydraulic conductance of the rhizosphere through an increase in root length. Such results remained to be tested at larger scale but could change our view about the mechanisms of species co-existence. Whereas it is sometimes assumed that mixture has positive effect due to root system segregation in space (Bello et al., 2019; Grossiord et al., 2018), we provide evidence that the hydraulic connection of the plant to the soil and the atmosphere can also be involved, without the need to call for a spatial segregation of the root systems.

Our results also challenge the way vegetation models represent drought stress. To date, the majority of process-based models assume that soil water deficit in the rooting zone drives the water status of the plant. However, we provide evidence that changes in hydraulic connection from the soil can make the plant behave independently from soil water status. Implementing such processes in larger scale vegetation models could help explain and predict co-existence between species and drought induced effect on forest community. Such modelling approach could be a step toward the development of tools allowing to design drought resilient mixture.

## Materials and Methods

### Seedlings and experimental design

The experiment was set up during the summer 2021. It consisted in applying a drought treatment (watering stop) to potted *P. halepensis* and *Q. ilex* trees grown in monoculture or in mixture while monitoring ecophysiological variables at 5 different dates. Seedlings of *P. halepensis and Q. ilex* (one- and two-years old respectively) of equivalent dimensions were repoted in January 2020. 90 trees of each species were planted in 12 L containers, including two individuals per pot, either in monoculture or in mixtures. The soil was composed of sand (∼20%) and organic matters. Half of the pots were equipped with a physical barrier made of acrylic fabric (with 30µm mesh) that precludes root colonization from one side to the other of the pot but allow water transfer between the two separated compartments. From 2019 to June 2021, saplings were grown at the National Forestry Office of France (ONF) nursery in Cadarache (Southeast of France) and were watered twice a week to field capacity and fertilized once a week. One month before the start of the experiment (June 2021), pots were brought on the campus of INRAe (Avignon, France) to acclimate in the experimental greenhouse. The greenhouse was equipped with air temperature, a humidity (HD 9817T1) and radiation loggers. It included an independent regulation of climate through aeration (opening of the glasses or forced ventilation in the compartment) and cooling (humidification of the air entering through a “coolLJ box”). These systems allowed regulating the environment of the greenhouse according to the defined settings. In addition, the sidewalls of the greenhouse have been whitewashed to homogenize the radiation and the temperature. The temperature was kept between 25 and 35 °C, relative humidity (RH) between 40 and 75%, and maximum diurnal photosynthetically active radiation (PAR) below 1000 µmol.m^-^². s^-1^ (SI appendix, Fig. S5).

During acclimation period, watering was applied as in the nursery. Among the initial batch of 90 pots, we selected 54 pots for which the two trees were alive and had reached a height between 40 and 60 cm with less than 10cm heights differences between the two trees. Pots were divided into two batches: a batch of 6 pots per composition (36 pots in total) that was assigned to the drought experiment, and a batch of 3 pots per treatment (18 pots in total) that was assigned to a control treatment in which trees were maintained watered all along the season (two times a week). All pots were monitored once a week, from July 26 to August 18, for soil water and water potential and ecophysiological variables (leaf water potentials, leaf gas exchange, pots water loss-described below). The day before the beginning of the experiment, at the end of the afternoon, all pots were watered at saturation and weighted.

### Tree water potentials

Water potential was estimated through leaf water potentials of all trees measured at predawn once a week across the experimental period..The evening before measurements, for each tree, one leaf (*Q. ilex*) or small twig (*P. halepensis*) was covered with an aluminium foil and placed in a ziplock plastic bag. In addition, to limit tree nocturnal transpiration and allow water potential equilibration between the tree and the soil (Rodriguez-Dominguez et al., 2022), trees were covered with a plastic bag and a piece of wet paper was included under the plastic bag. Samples were collected before sunrise, between 4 to 5 am, kept into the ziplock and immediately placed in a cooler for water potential measurement. The 108 measurements were done randomly in less than 4 hours following sampling, with a scholander pressure chamber (PMS model 1505 D). At the beginning of the experimentation, midday water potentials of tree were measured between 1 and 2 PM, following the same procedures as described above for predawn water potential (leaf or twig covered with an aluminium foil and placed in a ziplock plastic bag). There were used to parametrize the model.

### Tree leaf gas exchanges

Leaf level gas exchange was measured using two portable photosynthesis system (LI-6400XT) for all trees at all dates except the second one due to breakdown of the greenhouse system. Measurements were done between 11 am to 3 pm, period during which PAR in the green house is highest and stable (between 600 and 1000 µmol.m^-^². s^- 1^). Licor chamber conditions were set to keep close to the greenhouse while providing non-limiting conditions: PAR was set at 1000 µmol.m^-^². s^-1^, the block temperature was set at 25°C, flow rate and scrubbing were adjusted to maintain RH between 60 and 80%. The leaves were allowed to acclimate for at least 3 minutes in the chamber before measurement, to ensure gas exchange stability. For each leaf *(Q. ilex)* or needle bunch (*P. halepensis*), ten values were recorded during one minute and the average was used in the data analysis. After the measurement, the area of leaves or needles included in the chamber were cut and stored in a plastic bag inside a cooler. The day after, leaf area was measured to correct gas exchange computation with actual leaf area in the chamber. Samples were then dried during 48 hours at 70°C to estimate specific leaf area.

### Tree biomass and leaf area estimates

We estimated leaf area of each tree at the beginning and the end of the experiment using a method relying on profile photographs, proposed by (Michael and Parker, 2000). It is based on a calibrated relationship between the projected area of the tree profile and the foliage biomass estimated destructively. For each species, we first built a calibration relationship between numbers of tree pixels in profile photographs and the foliage biomass. For the calibration relationship, trees were selected to span the range of sizes encountered in the experiment. We sampled trees before the beginning of the drought experiment (June 2021), and after the experiment (September 2021) to consider potential changes in size or leaf area or angulation that could have occurred during the summer and influenced the relationship. For each tree, the profile surface projected area was estimated by photography. All the settings were made to ensure a constant reproduction ratio (i.e., constant dimensions of real object dimensions per pixel) among photographs. To obtain foliage dry mass, all trees used for this calibration were cut at the base of the stem after taking photographs. Tree parts were sorted to separate green foliage, dead foliage, and the rest which is almost entirely made of stems. Tree parts were then dried at 70°C for 3 days (leaves/ needles) or until there was no variation in dry mass (almost one week). The leaf area of each tree with the estimation of total foliage dry mass at a specific date and specific leaf area estimated on leaf gas exchange measurement samples.

At the end of the experiment and for droughted pots, the belowground part of each tree were uprooted. The rooting system was washed to separate the soil particles for the roots. The rooting system extension (maximal length and width) were measured using a ruler, with a millimeter resolution. The root system was then dried out at 70°C in an oven for at least 10 days, until there are no more weight variations, and the total dry mass was estimated.

### Soil water content and soil water potentials

Pots were weighted at each measurement dates in the morning (8am) and at the end of the measurement day (5pm). Soil water content was estimated at the pot level, by subtracting the total pot weight, performed at each measurement dates in the morning (ca. 8 am), the soil dry mass and the total fresh tree biomass. Soil water potential (Ψ_soil_) was then estimated at the pot level from the normalized soil water content of the pots (*W _norm_*) and water retention curves determined in the laboratory on soil samples (V= 6 cm^3^). The determination of the retention curve was made with the combination of suction table (Ψ_soil_ > -0.01 MPa), pressure plate (Ψ_soil_ > -1.5 MPa) and dew point hygrometer (WP4C, Decagon-Ψ_soil_ < -1.5 MPa) methods (Dane and Hopmans, 2002). Five soil sample replicates were used for each point of the retention curve and the gravimetric water content was determined from fresh and dry weight obtained after drying in an oven at 70°C (limit temperature to avoid organic matter degradation) for about one week. To perfectly match the data, two different retention curves were fitted. A first retention curve was fitted with two set of van-Genuchten relationships (van Genuchten, 1980) intersecting at a gravimetric water content of 0.116 g.g^-1^ (corresponding to Ψ_soil_ = -1.4 MPa). A second set of retention curve was fitted with only one van-Genuchten relationships (van Genuchten, 1980). The retention curves take the following form:

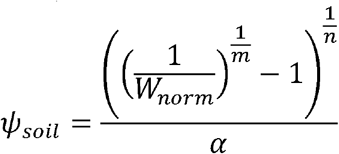

Where *m, n* and *α* are empirical parameters describing the typical sigmoidal shape of the function and *W*_*norm*_ is the normalized water content. Water potential was calculated from this fit using the gravimetric water content of pots estimated at each measurement dates. The parameters of the curves are provided in the SI appendix, Table S3.

The normalized water content was computed for each pot as:

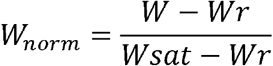

With *w* the soil mass of the pot at a given time, *Wr* the soil mass at residual water content. It was measured at the end of the experiment after drying the soil at 70°C. *W sat* is the saturated mass of the soil which was estimated from the first weight measurement of the experiment, after the pots were irrigated at saturation. *W* and *W sat* were computed by removing the mass of the tree and the pot to the total weight measured during the experiment (either from the balance or continuous load cell measurements). The total tree fresh was measured at the end of the experiment, by assuming that tree growth that could have occur during the experiment can be neglected due to the extreme drought experienced by the tree. *W _norm_* was not measured on the control (irrigated) pots.

### Soil resistivity measurement

Electrical resistivity of soil in pots was measured using electrical resistivity tomography (ERT). 4 pots (including one control) per modality (monoculture or mixture, with or without root separation system) were selected. On these pots, electrical resistivity was monitored with time over 2 radial planes, located at 1/3 and 2/3 of the pots’ height, by inserting 20 stainless steel screws (2cm long) equally spaced (3.9cm) along the column’s circumference. ERT measurement were done using an ABEM SAS 4000 resistivity meter connected to all these electrodes. All quadrupole combinations were used, including reciprocal measurements for assessing error and measurement quality. The resistivity measurements were taken before the start of the experiment (when the pot substrates were at field capacity), in the middle and at the end of the experiment. In the late dry situations, it was necessary to add a small amount of water at electrodes to enable soil-electrode electrical contact and resistivity measurements. Soil resistivity distribution at the two heights was obtained from the inversion of apparent resistivity using ResIPy software (Blanchy et al., 2020).

### Statistics

We evaluated globally the effect of species and measurement date and their interaction on the water potential of trees by using a linear mixed model. Then, for each species independently and root separation modalities (root separation or not), we assessed the effect of the pot composition (mixture or monoculture association) on tree water potentials by considering date, composition and their interaction as explanatory factors. As we did not find any significant differences between water potentials of monoculture with and without root separation for each species (SI appendix, Fig. S6), we decided to pool them for the analysis. We also test the differences between soil and water potentials of tree at each measurement dates using Student T test. Finally, we applied post-hoc Tuckey HSD tests to evaluate differences between pots modalities (composition and root separation modalities) for leaf area and gas exchanges variables (leaf conductance and transpiration, Fig. S7). All statistical analyses were performed with the R software (3,5,2, R Development Core Team 2018) with the package LME4 and *agricolae*.

### Model analysis using SurEau

We used the SurEau model to explore how the species composition in pots influenced soil and water potential dynamics during extreme drought (Cochard et al., 2021). SurEau is a soil-tree-atmosphere model that simulates water fluxes and stocks with the soil-tree atmosphere continuum by accounting for conductance, water potential gradient and capacitances. It is dedicated to model extreme drought and accounts for the processes occurring after the point of stomatal closure (*i.e.,* cuticular water losses and hydraulic conductance and water stocks losses due to xylem embolism). It is discretized in the soil layers and four tree compartments (roots, trunk, branches, and leaves) which are d each described by an apoplasmic and a symplasmic water volume. At each time step, the model computes leaf stomatal and cuticular transpiration as the product between leaf-to-air vapor pressure deficit and stomatal and cuticular conductance. Knowing the soil water content, soil water potential and hydraulic conductance are computed. This along with leaf stomatal and cuticular fluxes, can be used to compute tree water potential in the different tree compartments while accounting for the symplasmic capacitance and the hydraulic conductance losses due to xylem embolism. The resulting water potential is used to compute stomatal closure, water content, and xylem embolism. The model is driven by hourly climate data, tree traits (water pools dimensions, stomatal response to water potential, cuticular transpiration, capacitance, and vulnerability curve to cavitation (SI appendix, Table S4) and soil properties (volume and water retention curves). In the present study, the model was improved to include the possibility for two trees to absorb water in the same soil volume. In principle, two codes corresponding to two trees, parameterized for monoculture of *P. halepensis*, monoculture of *Q. ilex* or for mixture, were run in parallel.

To test the hypotheses presented in the introduction (illustrated in Fig. 1) and evaluate whether the model can also explain the patterns of soil and water potential dynamics found experimentally. First, we performed *benchmark* simulations, for monoculture and mixture, with default parameters, to reproduce the hypotheses presented in Figure 1. As explained above, the patterns of Figure 1 hold only under the assumptions that (i) the two individuals in the pots exploit the same soil water stock (physical coexistence of the roots) and are perfectly connected to the soil (large hydraulic conductance between the soil and the fine roots). In SurEau the flow of water between the soil and the roots is modelled using two different types of conductance in series, (i) the hydraulic conductance between the soil and the root surface (K_soil_) which depends on fine roots density and the soil water content (Martin-StPaul et al., 2017), and (ii) the hydraulic conductance between the root surface and the inner root (K_root_) which depends on the fine roots area (Ra) and fine roots conductivity (K_root_), both set constant by default (Cochard et al., 2021). The first hypothesis (root occupy the same volume) was fulfilled by setting the same quantity of fine roots in all three soil layers for the two individuals in the pot. The second hypothesis (equilibration between tree and soil water potential) was fulfilled by setting a fine roots length so that the soil hydraulic conductance is high enough for the night-time water potential equalled the soil water potential all along the drought range and before hydraulic failure (where water potential drops to minus infinity).

However, two different empirical results conflicted the expected patterns and we used the model to explain the divergences observed.

Firstly, *P. halepensis* showed that water potential can be higher than soil water potential during extreme drought under all composition monoculture and mixture, suggesting that this species can behave independently from the soil and maintain its water potential constant even if soil water potential decreases. Recent work suggests that disconductance between the soil and root can occur for some species (Cuneo et al., 2016; Duddek et al., 2022; North and Nobel, 1997). This can be represented in the model by decreasing the root conductance relative to water potential in the root. We thus implemented an equation relating the root hydraulic conductance to the root water potential (SI appendix, Fig. S2) for *P.halepensis* and realized simulation in mixture conditions with *Q. ilex* parametrized as in benchmark conditions. It appeared that this implementation led to an acceleration of hydraulic failure for *P.halepensis*. This is explained by the excessive water losses, that occur through the cuticle at this stage of water stress, and that cannot anymore be compensated by the supply from the root. Under the same mixture conditions, we therefore also tested whether a decrease in leaf cuticular conductance with relative water content (RWC, SI Appendix, Fig. S3), a phenomenon already observed on cut branches of *P. halepensis*, could explain -- alone or in combination with the reduction in root conductance -- the observed pattern.

Secondly, for *Q. ilex*, we noticed lower water stress under mixture linked to a change of the soil water potential (Ψ_soil_) *vs* plant predawn water potential (Ψ_pd_) relationship. Higher plant water potential for a given soil water potential was found under mixture compared to monoculture. Such pattern could be explained by an increase of the soil hydraulic conductance that, as explained above, can be related to the density of fine roots (*L*_*a*_, the length of fine roots per m² of soil). We thereby performed a sensitivity analysis on this trait under monoculture conditions to support that its variation could explain the empirical observation.

## Acknowledgments

Myriam Moreno was supported by the French Environment and Energy Management Agency (ADEME) in the form of a PhD scholarship. The experiment was funded by Agence Nationale pour la Recherche (ANR Hydrauleaks, MixForChange) and the Metaprogramme ACCAF Drought&Fire.

## Author Contributions

M.M., N.M. and H.C designed the research; M.M., G.S., N.M. C.D, P.F and R.D. performed research; H.C. and N.M. performed the model simulations; M.M analyzed the data, with the help of N.M, G.S, and CD; M.M and N.M wrote the paper, and all authors contributed to its review.

