## Supplemental materials for "Functional diversity reduces the risk of hydraulic failure in tree mixtures through hydraulic disconnection"

*Myriam Moreno (corresponding author).

Supporting Information Text

#### Overview of SurEau and its parametrization

SurEau(1) computes the plant water fluxes from the soil to the atmosphere, the water potential of the soil and of the different plant organs, and the level of xylem embolism. The soil is divided in 3 layers for which the variations of soil conductance and water potential are computed. The plant is divided into four simplified organs: roots, one trunk, one branch and one leaf canopy. Each plant compartment includes a *symplasm* and an *apoplasm* that is characterized by a water volume, and for which water potentials are computed considering the incoming and out coming flows as well as the contribution of capacitance and cuticular transpiration (or water leaks computed only from the symplasm to the air). For symplasmic compartment, the capacitance is derived from pressure volume curve (PV curve, (2)) equations whereas for apoplasmic compartment, a constant capacitance is used. The resistance between the different compartments is set from direct measurements or from allometry if not readily available. The code starts with the computation of transpiration (stomatal and cuticular) and leaf energy balance using hourly time steps climate data. The stomatal conductance is computed with the Jarvis model that includes responses to light, temperature and leaf water status. The CO_2_ effect was omitted in this version. Stomatal response to leaf water status is accounted by using a function of leaf turgidity derived from species-specific leaf pression volume curves. Therefore, complete stomatal closure is achieved at turgor loss point (Ψ_tlp_). Cuticular transpiration of the different symplasms is computed from cuticular conductance and vapour pressure deficit. The loss of conductance due to embolism formation is described by the xylem vulnerability curves for each apoplasmic compartment.

Input climate data (temperature, air humidity, atmospheric pressure, global radiation and wind speed) measured in the greenhouse were used at hourly time step. The water volumes of the different compartments and the resistance between the different compartments were set according to average species-specific measurements from the experiment (Table S4). The total resistance of the plant was computed using leaf level transpiration (derived from licor measurements), midday and predawn water potential and disaggregated using assumption about the distribution of resistance in the plant: most of resistance is located in root and leaves symplasm). More specifically, we distributed the resistance as follow(3,4) 20% of the resistance in the leaf symplasm, 20% in the leaf apoplasm, 8% in the branch apoplasm, 2% in the stem apoplasm, 10 % in the root apoplasm and 40% in the root symplasm. The radial symplasmic resistance was computed for each woody compartment (roots, trunk, branch) using the developed areas and a symplasmic resistivity of 1 (mmol.m-². s-^1^. MPa-^1^) for trunk and branches and 3.5 (mmol.m-².s-^1^.MPa-^1^) for roots. Note that the root symplasmic hydraulic conductance (K_root_) dictates the water fluxes between the soil and the inner part of the root. The fine (absorbing) root length is a key parameter in the model that allows to compute soil hydraulic conductance. This was computed using fine root area, which was set equal to the leaf area, and used to compute the root length, by assuming a fine root radius of 0.5 mm. For the sake of simplicity, potential segmentation of xylem vulnerability was omitted and the same vulnerability curve to cavitation was used for all compartments of the same species. Similarly, the same species-specific leaf PV curve was used to compute the contribution of symplasmic capacitance to water flow of all the symplasmic compartments. The “benchmark simulations” were done for both monocultures and mixture using the parameters in Table S4.

#### Sensitivity tests and hypothesis about K_root_, g_cuti_ and root length

Testing the “isolation effect”: Can we explain the relatively constant water potential of *P. halepensis* with variable root hydraulic conductance (K_root_) and cuticular conductance (g_cuti_)?

To test the hypothesis about the possible influence of a variable symplasmic root hydraulic conductance (K_root_) and cuticular conductance to explain the drought avoiderity of water potential (constant water potential) in *P. halepensis,* we included additional equations in the model.

First, we implement a variable K_root_ by assuming a variable gap fraction in the root cortex:

$$K_{root}=k\_Root\_Symp0 * Root\_Area\_fi*\frac{(100-Cortex\_Gap)}{100}$$

With $k\_Root\_Symp0$ the initial hydraulic symplasmic conductivity, $Root\_Area\_fi$ the fine root area and $Cortex\_Gap$ the proportion of gap in the root cortex which is computed as sigmoidal function of the root symplasmic potential $P\_Root\_Symp$

$$Cortex\_Gap=\frac{100}{(1+exp(K\_VAR\_P2/25*(P\_Root\_Symp-K\_VAR\_P1)))}$$

with K_VAR_P1 the water potential causing 50% of cortex gap and K_VAR_P2 the slope at the inflexion point of the sigmoidal function.

To test the hypothesis that a variable cuticular conductance (g_cuti_) can combine with a variable K_root_ to maintain a relatively constant plant water potential for the *P. halepensis* under severe water deficit, we added a function to reduce the cuticular conductance. Precise measurements of cuticular conductance obtained using a drought-box(5) shows a reduction of g_cuti_ according to the relative water content of samples (see Figure S3 A just below). Hence, in the model we assumed that after turgor loss point, g_cuti_ at reference temperature (20°C) decrease with the relative water content of the leaves:

$$if (RWC\_leaf<RWC\_tlp)$$

$$g_{\mathrm{cuti}}=g_{\mathrm{cuti}_{20}}*\left( 1-\left( \mathrm{RWC}_{\mathrm{tlp}}-\mathrm{RWC}_{\mathrm{leaf}} \right)*\mathrm{RWC}_{\mathrm{sens}} \right)$$

$$\mathrm{else}g_{\mathrm{cuti}}= g_{cuti20}$$

With $RWC\_leaf$ the leaf symplasmic relative water content, $\mathrm{RWC}_{\mathrm{tlp}}$ the leaf relative water content at turgor loss point, $\mathrm{RWC}_{\mathrm{sens}}$ the sensitivity of $g_{\mathrm{cuti}}$ to relative water content, $g_{cuti20}$ the leaf cuticular conductance at 20°C.

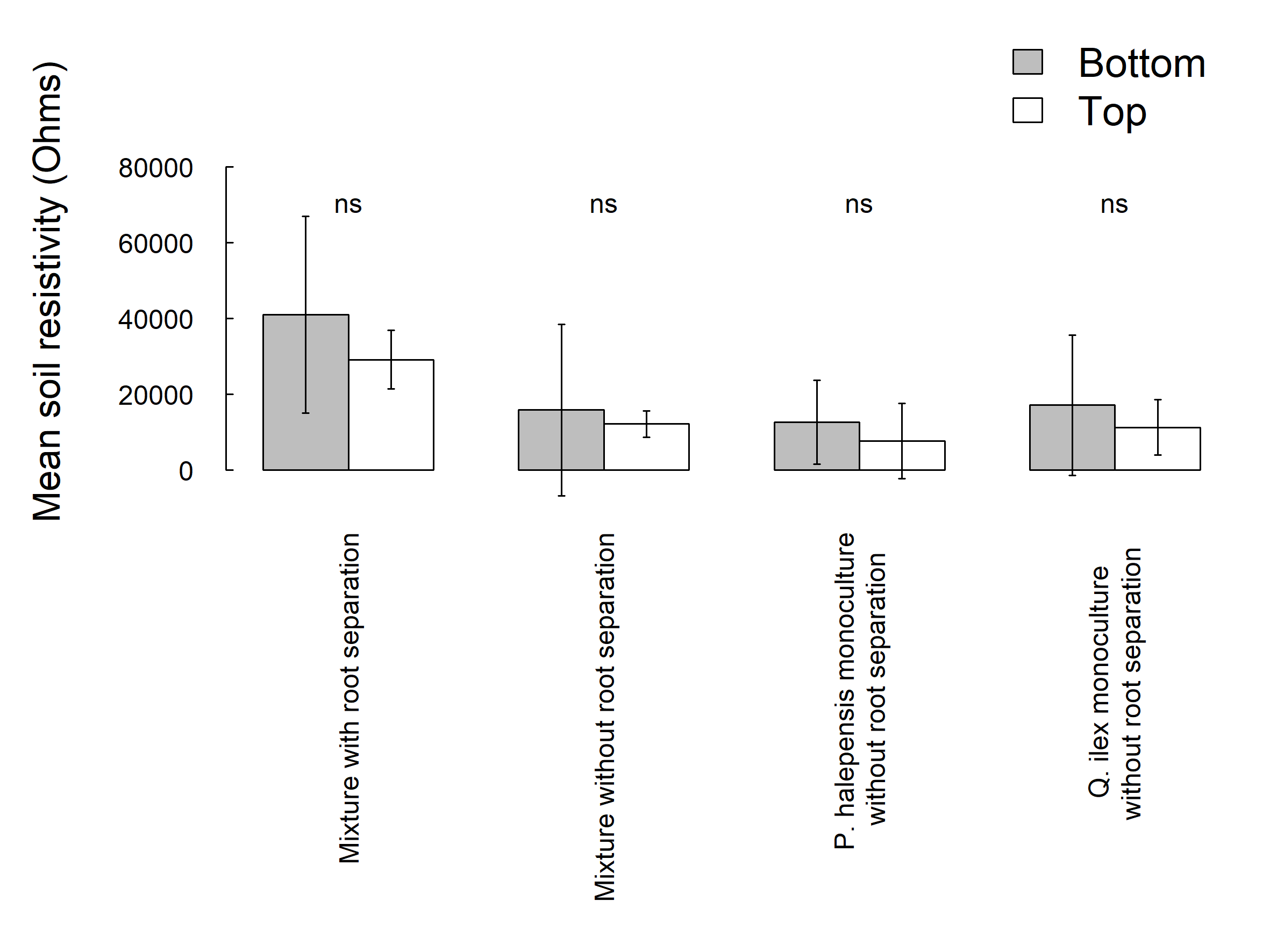

Fig. S1. Mean resistivity of the top and bottom profiles of pots (1/3 and 2/3 of the height of pot) at the end of the experiment according to pot modality. Non-significant differences were found between levels of measurement (top and bottom) in each kind of pot modality. Bars represent standard deviation. N=3 per pot modalities.

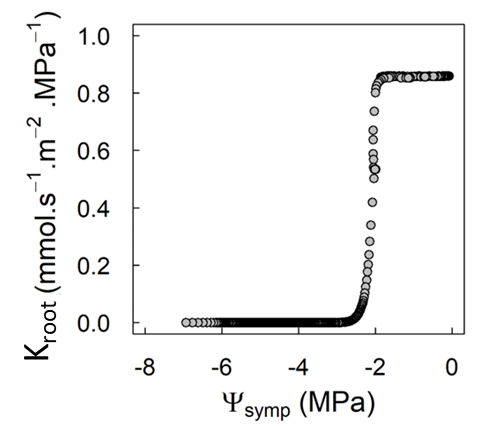

**Fig. S2**. Variation of root conductance (K_root_) according to root symplasmic water potential.

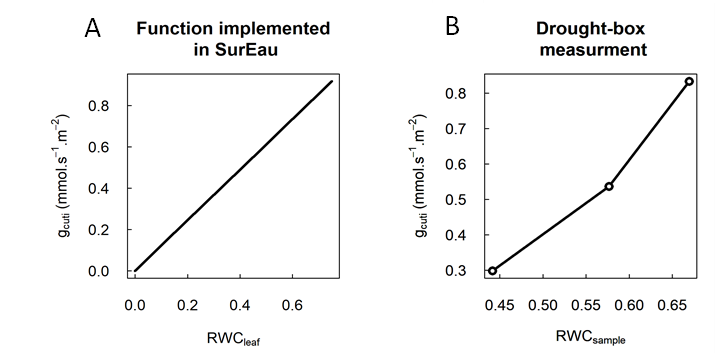
Fig. S3. Variation of g_cuti_ according to relative water content. A. Function implemented in the model. B. g_cuti_ measurements for a given sample of shoot dried under the drought-box (5).

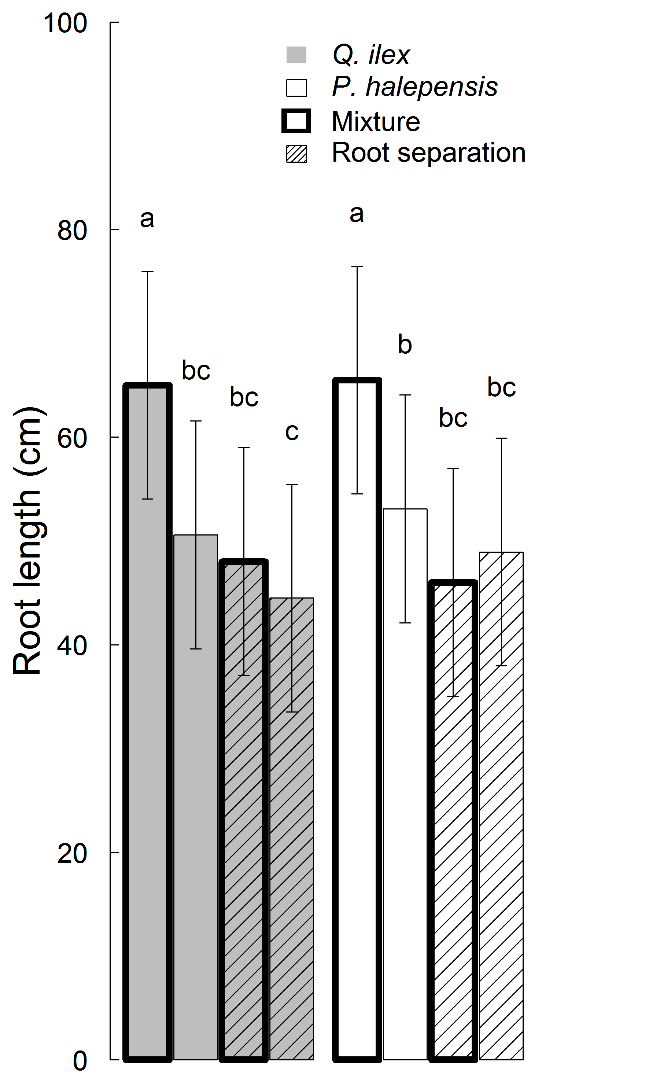

Fig. S4. Average root length of *Q. ilex* (grey bars) and *P. halepensis* (white bars) for the different pot composition modalities. Mixtures are represented with bold border and root separation with hatched bars. Root lengths were measured at the end of the experiments, after uprooting the pots carefully. For each species, N= 12 for monoculture and N= 6 for mixture.

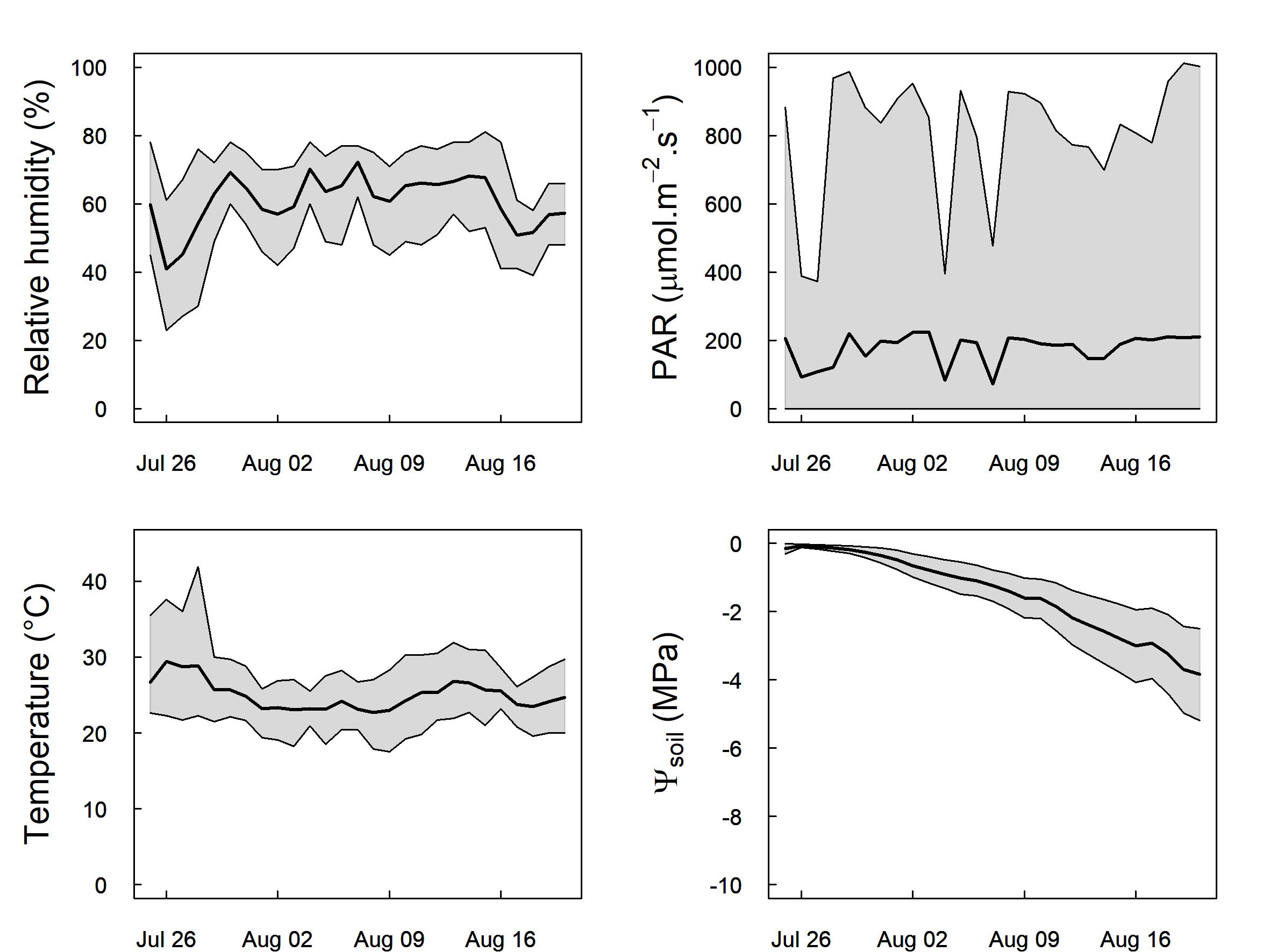

Fig. S5. Meteorological variables recorded in the greenhouse compartment during the experimentation: Relative humidity (%), Photosynthetic Active Radiation (PAR), Temperature (°C). For these three plots, the bold black line represents the average value and the grey interval the maximum and minimum value recorded. The measurements were recorded at a half-hourly timescale during the experimentation. The last plot represents average water potentials of the droughted (bold black line) pots (N= 36) and standard deviation (grey interval).

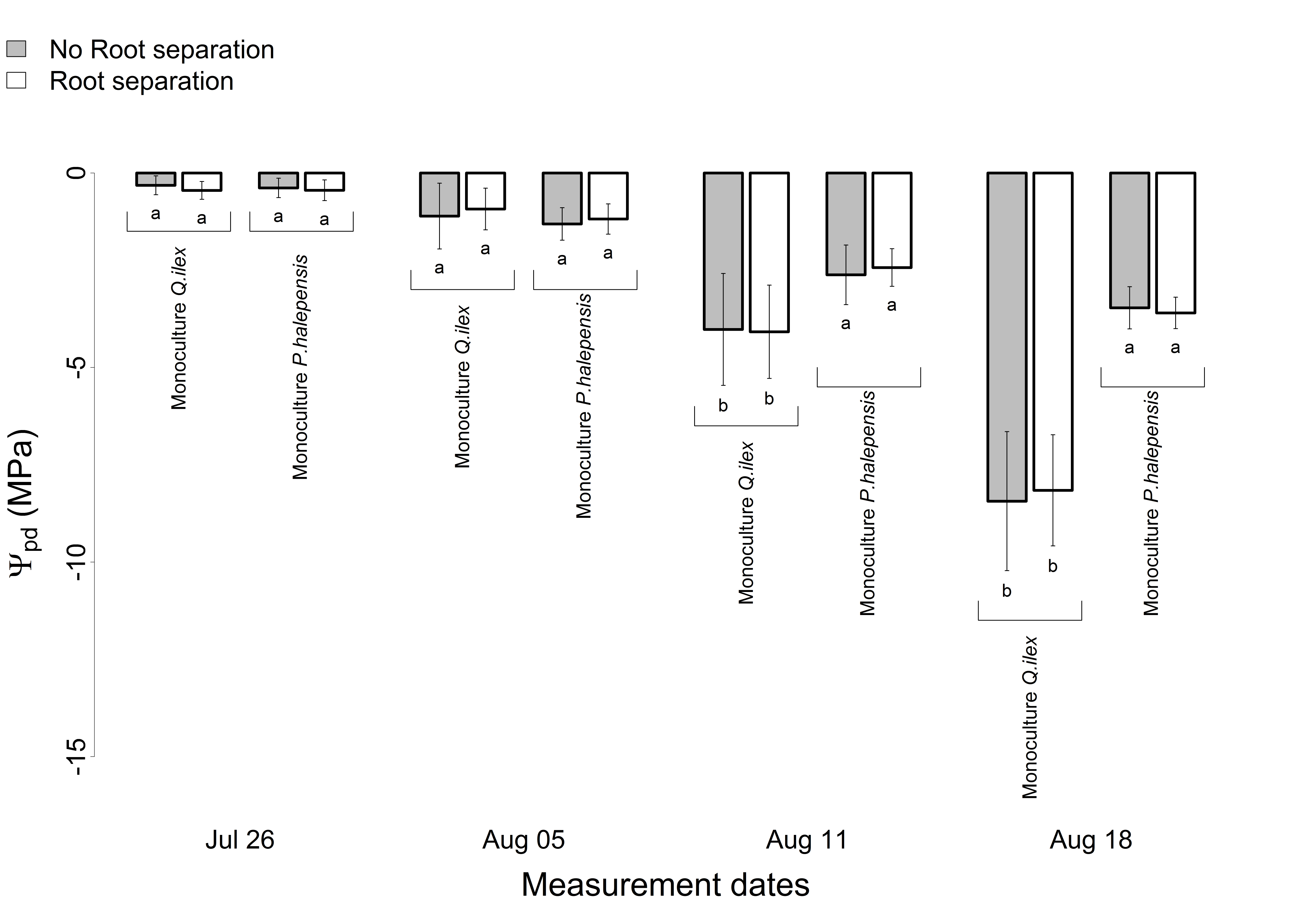

Fig. S6. Average and standard deviation of plant predawn water potentials in monoculture. Grey and white bars represent respectively pots without and with root separation. Letters represent the results of the Tukey post-hoc test comparing predawn water potential of the different pot modality at each measurement date. Per date, N= 6 in both monoculture with and without root separation.

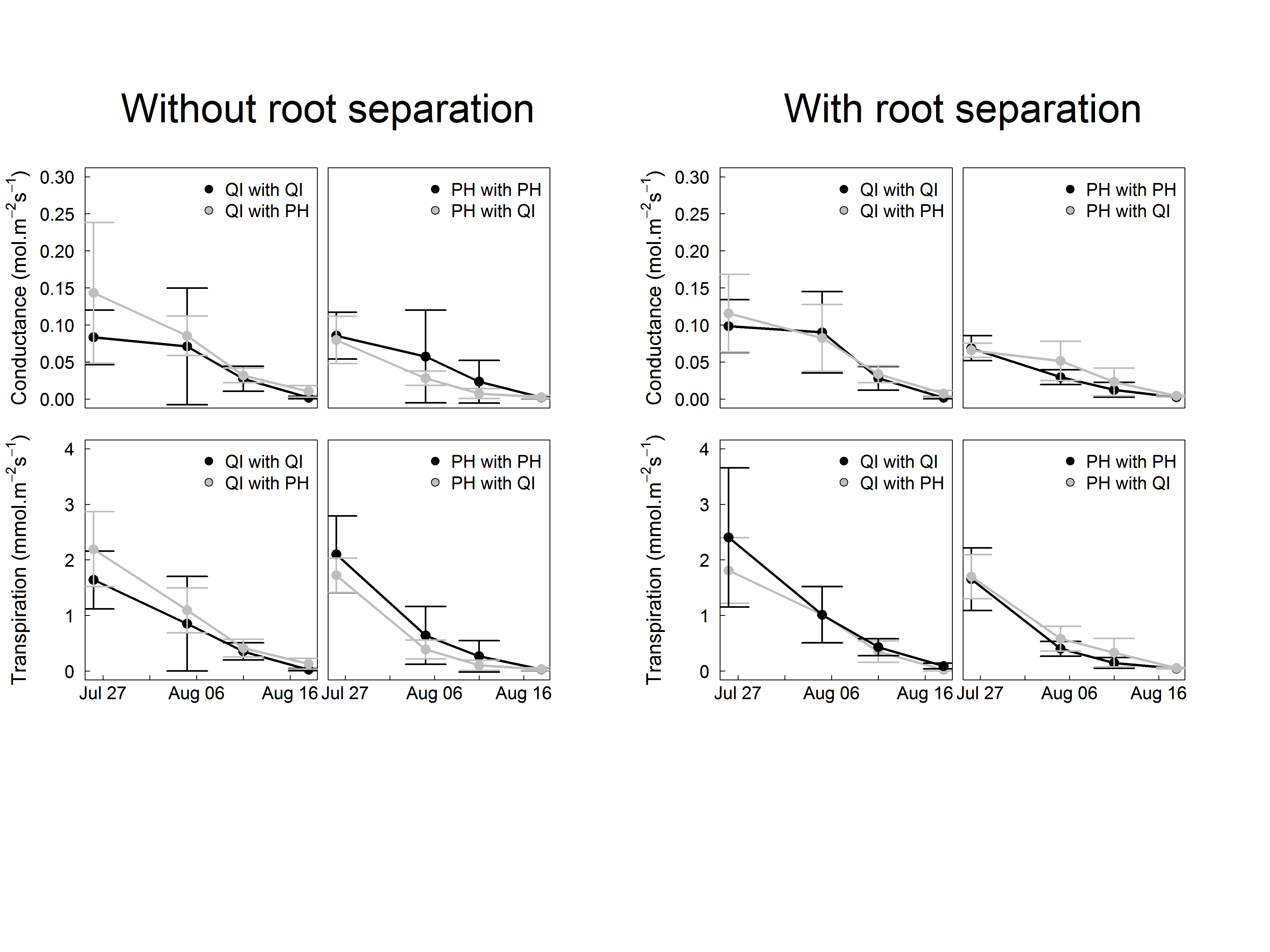

Fig. S7. Dynamics of the transpiration (mmol.m^-2^. s^-1^) and conductance (mol.m^-2^. s^-1^) at the leaf scale for *Q. ilex* (QI) in monoculture (black dots) or mixture (grey dots) and *P. halepensis* (PH) in monoculture (black dots) and mixture (grey dots) without (left panel) or with (right panel) root separation. Intervals represents standard deviation. In these plots, as Tukey’s HSD Post-doc tests report non-significant differences at each measurements date, no letters were added at the top of the bars. N > 3 for each measurement date excepting the last one where it was hard to make measurement.

**Table S1.** Analysis of variance of the mixed model for tree water potential (measured at predawn) to test date and species effect. The model includes a Species, a Date effect and Species*Date effect. An individual random effect has been included.

| **Factors** | **Sum Sq** | **Mean Sq** | **NumDF** | **DenDF** | **F value** | **Pr (>F)** | **Significance** |
| --- | --- | --- | --- | --- | --- | --- | --- |
| **Species** | 38.09 | 38.09 | 1 | 68.333 | 84.062 | 1.606e-13 | *** |
| **Date** | 1183.69 | 394.56 | 3 | 201.048 | 870.755 | < 2.2e-16 | *** |
| **Species:Date** | 225.97 | 75.32 | 3 | 201.048 | 166.230 | < 2.2e-16 | *** |

**Table S2.** Analysis of variance of the mixed models to test the composition effect (monoculture vs mixture), per species and type of pot (with or without root separation), for tree water potential (measured at predawn). The model includes a fixed effect date, a fixed effect mixture treatment and their interaction. For all model a random effect individual has been included.

| **Species and pot modalities** | **Factors** | **Sum Sq** | **Mean Sq** | **Num**  **DF** | **Den**  **DF** | **F value** | **Pr (>F)** | **Significance** |
| --- | --- | --- | --- | --- | --- | --- | --- | --- |
| ***Q. ilex* - pots without root separation** | **Date** | 470.2 | 156.731 | 3 | 47.440 | 219.4772 | < 2.2e-16 | *** |
|  | **Mixture vs Monoculture** | 2.09 | 2.085 | 1 | 16.319 | 2.9203 | 0.106417 |  |
|  | **Date:Mixture** | 11.08 | 3.694 | 3 | 47.440 | 5.1725 | 0.003566 | ** |
| ***Q. ilex* - pots with root separation** | **Date** | 578.38 | 192.794 | 3 | 48 2 | 97.1819 < | 2e-16 | *** |
|  | **Mixture vs Monoculture** | 0.26 | 0.260 | 1 | 16 | 0.4005 0 | .5358 |  |
|  | **Date:Mixture** | 1.31 | 0.437 | 3 | 48 | 0.6732 0 | .5727 |  |
| ***P. halepensis* - pots without root separation** | **Date** | 90.045 | 30.0151 | 3 | 42 1 | 88.7433 < | 2e-16 | *** |
|  | **Mixture vs Monoculture** | 0.072 | 0.0718 | 1 | 14 | 0.4516 0 | .5125 |  |
|  | **Date:Mixture** | 0.222 | 0.0739 | 3 | 42 | 0.4645 0 | .7086 |  |
| ***P. halepensis* - pots with root separation** | **Date** | 84.333 | 28.1111 | 3 | 46.184 | 242.9660 | <2e-16 | *** |
|  | **Mixture vs Monoculture** | 0.307 | 0.3072 | 1 | 16.430 | 2.6553 | 0.1222 |  |
|  | **Date:Mixture** | 0.257 | 0.0856 | 3 | 46.184 | 0.7399 | 0.5337 |  |

**Table S3.** Fitted parameters of the van Genuchten water retention curves to compute pot water potential (see M&M in the main text). In order to perfectly fit to the data, two curves were adjusted corresponding to two range of water content.

|  | **First water retention curves**  **(for W > 0.1214 g/g)** | **Second water retention curves**  **(for W < 0.1214 g/g)** |
| --- | --- | --- |
| **Wsat (g/g)** | 0,99528743 | 1,1 |
| **Wr (g/g)** | 0,06996382 | 0 |
| **alpha (1/bar)** | 374,7136575 | 2,21.10^1^ |
| **n** | 1,341822501 | 1,39185236 |
| **m** | 0,254744946 | 0,28153299 |

**Table S4. SurEau parameters used according to species.**

|  | **Leaf area (m²)** | **LMA (g.m^-2^)** | **Succulence**  **(gH2O.^m-2^)** | **E Licor**  **(mmol.m-².s^-1^)** | **gs_max_**  **(mmol.m-².s^-1^)** |
| --- | --- | --- | --- | --- | --- |
| ***P. halepensis*** | 0,19 | 164 | 300 | 2,2 | 185 |
| ***Q. ilex*** | 0,12 | 149 | 145 | 1,9 | 164 |
| **Source** | This study | This study | Unpublished data | This study | This study |
|  | **Ψ midday (MPa)** | **Ψ predawn (MPa)** | **k_plant_**  **(Leaf_specific, mmol.m-².s^-1^.MPa^-1^)** | **K_plant_**  **(mmol.s^-1.^MPa^-1^)** | **g_cuti_ 20°C (mmol.m-².s^-1^)** |
| ***P. halepensis*** | -1,06 | -0,2 | 2,1 | 0,35 | 1,1 |
| ***Q. ilex*** | -1,13 | -0,2 | 1,68 | 0,235 | 2,38 |
| ***Source*** | This study | This study | This study | This study | Unpublished data |
|  | **Ψgs_90 (MPa)** | **P50_Xylem (MPa)** | **Ψ_tlp_ (MPa)** | **π100 (MPa)** | ε **(MPa)** |
| ***P. halepensis*** | -2 | -5 | -2,1 | -1,6 | 7 |
| ***Q. ilex*** | -3,3 | -7 | -4 | -3 | 12 |
| **Source** | Unpublished data | Unpublished data | Unpublished data | Unpublished data | Unpublished data |
|  | **Apoplasmic Fraction**  **(unit-less)** | **Moisture content in wood (gH_2_0/gMS *100)** | **Moisture content in leaves**  **(gH20/gMS *100)** |  |  |
| ***P. halepensis*** | 0,6 | 130 | 165 |  |  |
| ***Q. ilex*** | 0,4 | 80 | 80 |  |  |
| **Source** | Unpublished data | This study | This study |  |  |
